# Multiscale model of integrin adhesion assembly

**DOI:** 10.1101/542266

**Authors:** Tamara Carla Bidone, Austin V. Skeeters, Patrick W. Oakes, Gregory A. Voth

## Abstract

The ability of adherent cells to form adhesions is critical to several phases of their physiology. The assembly of adhesions is mediated by several types of integrins. These integrins differ in physical properties, including rate of diffusion on the plasma membrane, rapidity of changing conformation from bent to extended, affinity for extracellular matrix ligands, and lifetimes of their ligand-bound states. However, the way in which nanoscale physical properties of integrins ensure proper adhesion assembly remains elusive. We observe experimentally that both β-1 and β-3 integrins localize in nascent adhesions at the cell leading edge. In order to understand how different nanoscale parameters of β-1 and β-3 integrins mediate proper adhesion assembly, we therefore develop a coarse-grained computational model. Results from the model demonstrate that morphology and distribution of nascent adhesions depend on ligand binding affinity and strength of pairwise interactions. Organization of nascent adhesions depends on the relative amounts of integrins with different bond kinetics. Moreover, the model shows that the architecture of an actin filament network does not perturb the total amount of integrin clustering and ligand binding; however, only bundled actin architectures favor adhesion stability and ultimately maturation. Together, our results support the view that cells can finely tune the expression of different integrin types to determine both structural and dynamic properties of adhesions.

**Author summary:** Integrin-mediated cell adhesions to the extracellular environment contribute to various cell activities and provide cells with vital environmental cues. Cell adhesions are complex structures that emerge from a number of molecular and macromolecular interactions between integrins and cytoplasmic proteins, between integrins and extracellular ligands, and between integrins themselves. How the combination of these interactions regulate adhesions formation remains poorly understood because of limitations in experimental approaches and numerical methods. Here, we develop a multiscale model of adhesion assembly that treats individual integrins and elements from both the cytoplasm and the extracellular environment as single coarse-grained (CG) point particles, thus simplifying the description of the main macromolecular components of adhesions. The CG model implements sequential interactions and dependencies between the components and ultimately allows one to characterize various regimes of adhesions formation based on experimentally detected parameters. The results reconcile a number of independent experimental observations and provide important insights into the molecular basis of adhesion assembly from various integrin types.

## Introduction

As the linker between cytoskeletal adhesion proteins and extracellular matrix ligands, integrins play a vital role in the formation of adhesions and profoundly influence different phases of cell physiology, such as spreading, differentiation, changes in shape, migration and stiffness sensing, [1–5].

Integrins are large heterodimeric receptors, with a globular headpiece projecting more than 20 nm from the cell membrane, two transmembrane helices, and two short cytoplasmic tails that bind cytoskeleton adhesion proteins (see Fig 1A). In order to form adhesions, integrins undergo lateral diffusion on the cell membrane, switch conformation from bent to extended, and change chemical affinity for extracellular matrix (ECM) ligands, *E_IL_*. Integrins also assemble laterally, owing to interactions with talin [6,7], kindlin [8], or glycocalyx [9], and can grow nascent adhesions into mature adhesions [10–12]. Integrin diffusion, activation, ligand binding, and clustering occur at the individual protein scale, but their effects can also be reflected on the cellular scale, resulting in a multiscale biological process. Simulations of adhesion assembly based on all-atom approaches are too detailed and computationally demanding to capture adhesion formation from multiple integrins. Instead, highly CG, approaches based on Brownian Dynamics can condense the description of individual proteins into a few interacting CG “beads” that can recapitulate the emergent dynamics of complex biological systems from its individual components (see, e.g., Refs [13–17] for the example of cytoskeleton networks).

**Fig 1.**
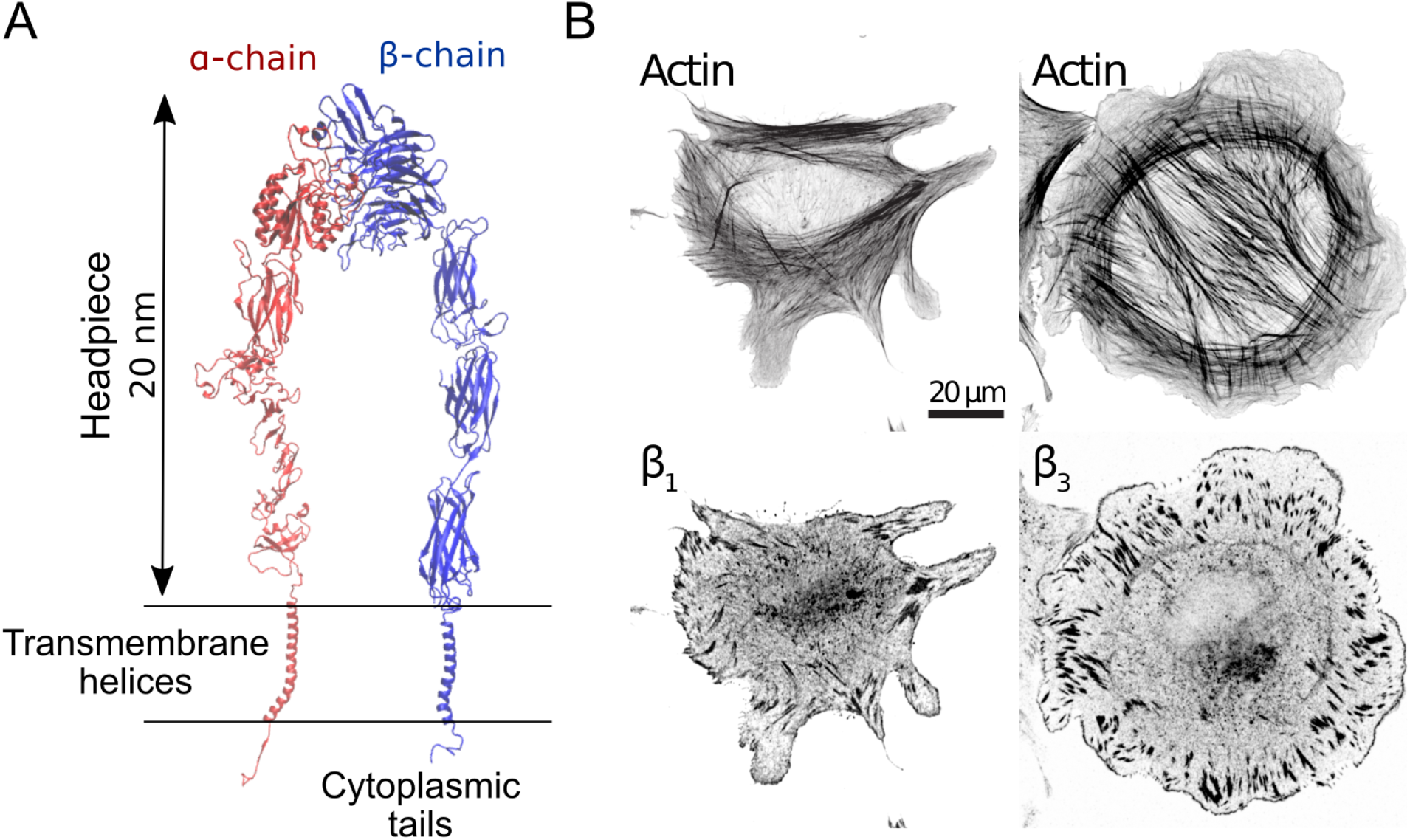
Both β-1 and β-3 integrins localize at the cell periphery, where nascent adhesions assemble. (A) Cartoon representation of active, fully extended *α_IIB_β*_3_integrin, which is closely related to *α_ν_β*_3_ [75]. (B) Representative images of human foreskin fibroblasts (HFF) fixed after 60 minutes of spreading on fibronectin coated glass coverslips and immunostained for actin and either β-1 or β-3 integrins.

Nascent adhesions are complex biological systems that form near the leading edge of protruding cells, appearing as spots of about 0.1 μm in diameter, with lifetimes of 2-10 min (Fig 1B) [18–22]. Unfortunately, the small size and short lifetime of nascent adhesions have made it challenging to study them experimentally. Among 24 different integrin isoforms, the *α_ν_β*_3_ and *α*_5_*β*_1_ integrins, have important, but potentially separate roles in the assembly of adhesions and the physiology of many cell types [21,23,32–34,24–31]. Nanoscale differences in physical properties between *α_ν_β_3_* and *α*_5_*β*_1_ integrins can determine how nascent adhesions assemble [35], their organization [36–38], transmitted traction [30,39] and lifetime [40], on account of their different properties. For example, it has been reported that the rate of integrin activation, *k_a_*, determines the number of integrins per adhesion [22,41], while lateral clustering, or avidity, *E_II_*, increases the size of individual adhesions [22,42–44].

Single-protein tracking experiments combined with super-resolution microscopy and computational methods have helped extract physical properties of two integrin types: For example, ß-1 and ß-3 integrins were found to have diffusion coefficients of 0.1 and 0.3 μm^2^/s, respectively [45]. ß-1 integrins also maintain their active conformation longer than ß-3 integrins. Free-energy energy differences between active and inactive states revealed activation rates for ß-3 integrins about 10-fold higher that ß-1 integrins [46,47]. The intrinsic ligand binding affinity, *E_IL_*, for soluble fibronectin is about 10-50 fold higher for ß-1 than ß-3 integrins [30], spanning an overall range for the two integrins of 3-9 *k_B_T* [48]. ß-1 integrins display a catch bond and adhesion strength-reinforcing behavior and are stationary within adhesions [45,49–51]. ß-3 integrins, on the other hand, rapidly transit from closed to open conformations, break their bonds from ligands more easily under modicum tensions, and undergo rearward movements within adhesions [45,52,53]. How these differences in diffusion, rate of activation, ligand binding affinity, and bond dynamics reflect on the assembly of nascent adhesions and on the probability of adhesion maturation remains elusive.

In this paper, we introduce a highly CG model of adhesion formation, based on Brownian Dynamics (Fig 2), in order to study how nanoscale physical properties of integrins determine adhesion morphology and stress transmission. The CG model treats individual integrins as point particles within an implicit cell membrane and includes actin filaments as explicit semiflexible polymers (Fig 2A). By incorporating nanoscale physical properties of individual integrins, sequential interactions and feedback mechanisms between integrin, ligands and actin filaments (Fig 2B-D), the model is used to characterize the formation of micrometer-size adhesions at the cell periphery in a multiscale fashion. Our calculations show that integrins with high *E_IL_* and enhanced bond lifetimes, such as ß-1 integrins, facilitate ligand binding, transmission of traction stress, and engagement of actin networks. By contrast, integrins with low *E_IL_* and lower ligand bond lifetimes, such as ß-3 integrins, are correlated with clustering, repeated cycles of diffusion and immobilization, and weak engagement of actin filaments. The architecture of actin filaments does not impact the amount of ligand binding and integrin clustering, but determines the probability of adhesions maturation, consistent with previous experimental findings [54]. Collectively, our data reveal important insights into adhesions assembly that are currently very challenging to obtain experimentally. The data supports the general view that cells, by controlling physical nanoscale properties of integrins via expression of specific types, can regulate structural, dynamical, and mechanical properties of adhesions.

**Fig 2.**
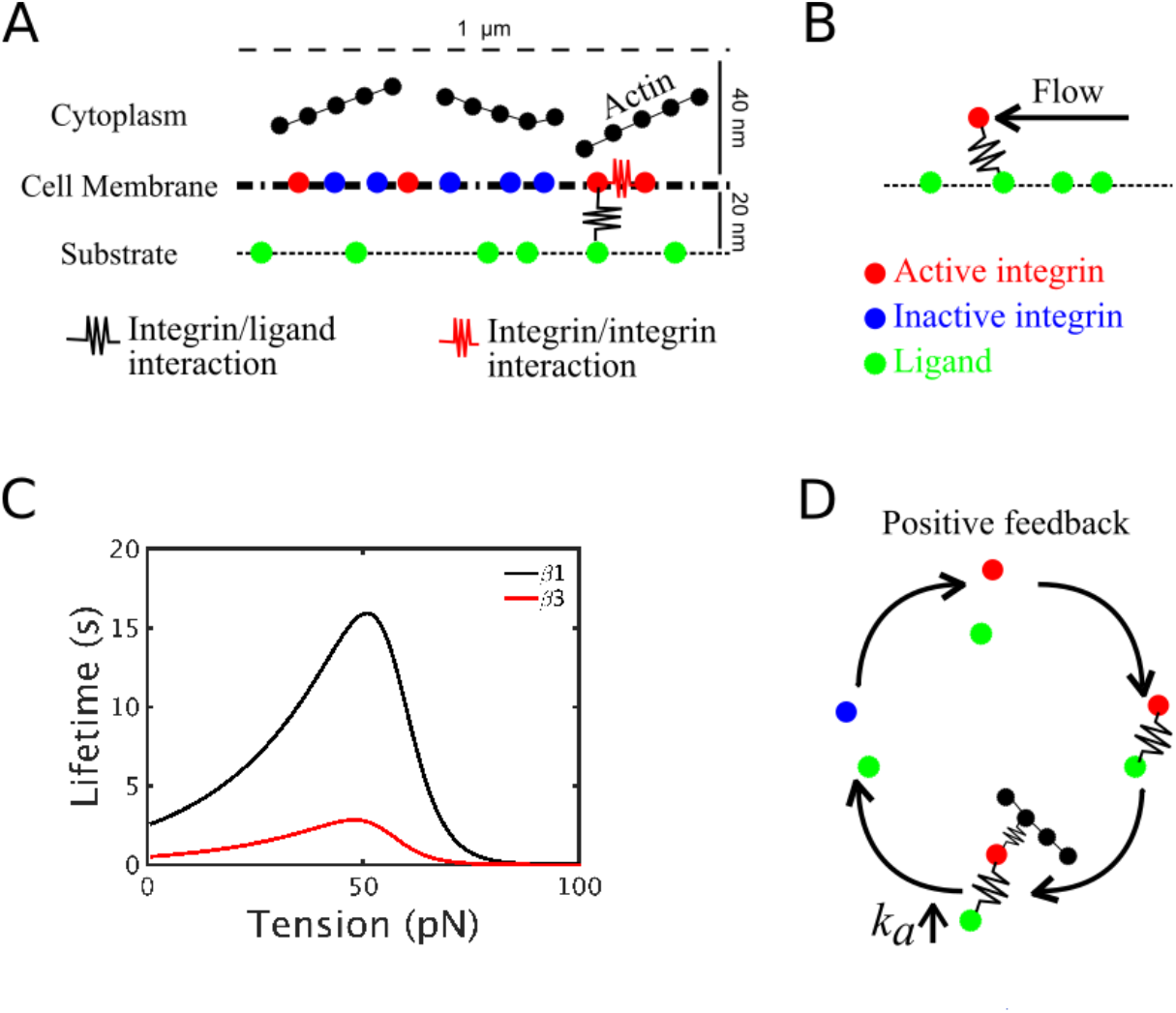
Schematic illustration of the model system. (A) Side view of the computational domain, where single-point, two-state integrin particles diffuse and assemble laterally. Upon activation (from blue to red), integrins can establish interactions (black spring) with ligands (green particles) and other active integrins (red springs). (B) Schematics of the system with actin flow: a force mimicking actin flow is exerted on ligand-bound integrins, parallel to the substrate, and builds tension on the integrin/ligand bond. (C) Lifetime versus tension curves of catch bonds used to mimic β-1 (black) and β-3 (red) integrins. (D) Schematic illustration of the positive feedback between actin filament binding and integrin activation. Once an integrin particle is bound to a ligand, it can establish interactions with actin. Upon binding actin, its activation rate increases. Upon deactivating and unbinding, its propensity to become active again increases, leading to increased probability of binding new ligands and actin filaments. This results in a positive feedback between ligand binding and engaging the actin cytoskeleton.

## Results

### Both β-1 and β-3 integrins localize in nascent and mature adhesions

Motivated by our recent work on integrin catch-bonds regulating cellular stiffness sensing [5], we sought to investigate how interactions between different integrins could affect adhesion formation. We stained Human Foreskin Fibroblasts (HFF) for actin and either β-1 or β-3 integrins. We found that both types of integrins localize in nascent adhesions at the cell leading edge and in mature adhesions at the end of actin stress fibers (Fig 1B). This suggests that potential interactions between the different adhesion populations could be important during adhesion formation. To address this question, we then developed a computational model to investigate how the nanoscale properties of different integrins affect adhesion formation and stability.

### Integrin clustering decreases with ligand binding affinity

Since β-1 and β-3 integrins differ in ligand binding affinity, *E_IL_*, and strength of pairwise interactions, *E_II_* [46,47], we use the CG computational model to test how variations in *E_II_* and *E_IL_* impact adhesion assembly in terms of the amount of integrin clustering, ligand binding, and spatial arrangement of adhesions. Different morphological arrangements of integrin adhesions are detected (Fig 3A-C). For high *E_II_* and low *E_IL_*, clustering is promoted (Fig 3D), but only a few integrins are bound to ligands (Fig 3E), resulting in few large integrin clusters (Fig 3A). Conversely, for low *E_II_* and high *E_IL_*, only a few integrins cluster (Fig 3D) while ligand-binding is promoted (Fig 3E), resulting in many ligand-bound integrins and few small integrin clusters (Fig 3B). When *E_II_* and *E_IL_* have intermediate values, a mix of big clusters of integrins that are weakly bound to the substrate and smaller, ligand-bound clusters co-exist (Fig 3C). By systematically varying *E_II_* and *E_IL_*, morphological regions differing in size and number of ligand-bound integrins versus clusters were precisely identified. A region of few large clusters exists for *E_II_* >3 *k_B_T* and *E_IL_* < 3 *k_B_T*; a region of many small adhesions exists for *E_II_* < 3 *k_B_T* and *E_IL_* > 3 *k_B_T*; the rest of the parameter space shows co-existence of intermediate-size clusters and ligand-bound integrins (Fig 3D-E). The fraction of ligand-bound integrins increases with *E_IL_* and is independent from *E_II_*. (Fig 3E). By contrast, clustering is not independent from *E_IL_* and is promoted when *E_IL_* is low (Fig 3D). In the model, when active, integrins can bind free ligands and cluster, when in close proximity of a ligand or another active integrin, respectively. Since the number of ligands is higher than the number of integrins, the probability for an integrin to find a free ligand is higher than that of finding an active integrin. Therefore, clustering increases less with *E_II_* when *E_IL_* is high than when *E_IL_* is low (Fig 3D). This indicates that integrin clustering and ligand binding are competing mechanisms.

**Fig 3.**
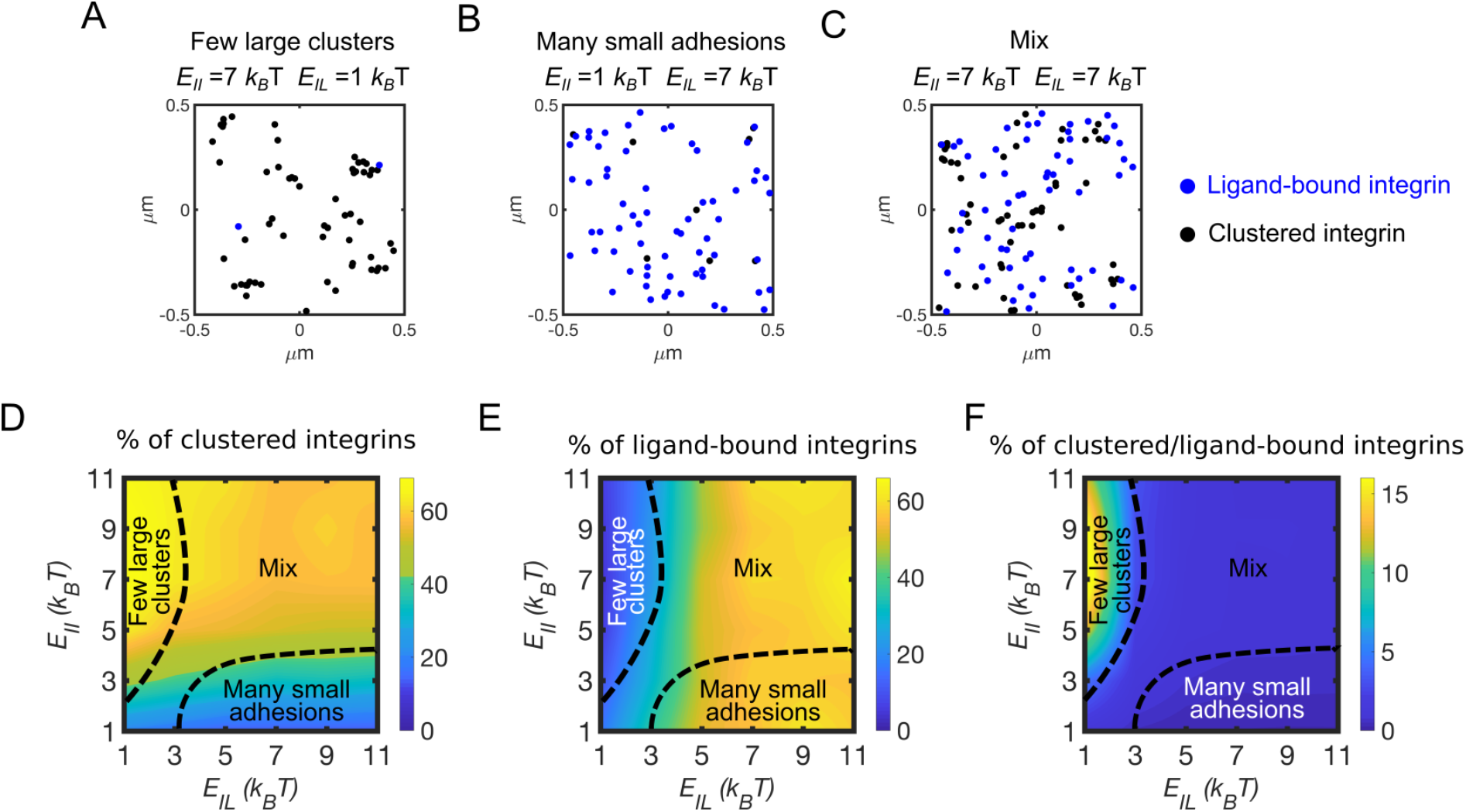
Integrin affinity and avidity determine clustering and ligand binding. (A) Configuration of clustered integrins (black circles) and ligand-bound integrins (blue circles) at a time point between 80-100 s of simulations, using *E_II_* = 7 *k_B_T* and *E_IL_* = 1 *k_B_T*. (B) Configuration of clustered integrins (black circles) and ligand-bound integrins (blue circles) at a time point between 80-100 s of simulations, using *E_II_* = 1 *k_B_T* and *E_IL_* = 7 *k_B_T*. (C) Configuration of clustered integrins (black circles) and ligand-bound integrins (blue circles) at a time point between 80-100 s of simulations, using *E_II_* = 7 *k_B_T* and *E_IL_* = 7 *k_B_T*. (D) Average percentage of clustered integrins relative to total integrins, by varying *E_IL_* and *E_II_*. (E) Average percentage of ligand-bound integrins relative to total integrins, by varying *E_IL_* and *E_II_*. (F) Fraction between clustered and ligand-bound integrins, by varying *E_IL_* and *E_II_*. This indicates the amount of clustered integrins per ligand-bound integrin. All data are computed between 100-130 s of simulations, from four independent runs.

Together, our results show that different arrangements of nascent adhesions can be achieved depending on *E_II_* and *E_IL_*. When we use high *E_II_* and low *E_IL_*, as for β-3 integrins, clustering is enhanced, and ligand binding reduced; when we use high *E_IL_* and low *E_II_*, as for β-1 integrins, clustering is reduced, and ligand-binding promoted. Thus, the competition between clustering and ligand binding can be determined by the integrin type. However, β-1 and β-3 integrins also differ in their rates of activation, which can lead to differences in this competition, by promoting clustering at high *E_IL_*. Therefore, we next aimed to understand how activation rates, combined with variations in *E_II_* and *E_IL_*, impact clustering and ligand binding.

### Rate of integrin activation increases clustering and ligand binding

Competition between integrin clustering and ligand binding can be determined by the difference in activation rate between β-1 and β-3 integrins. By varying *k_a_* from 0.005 s^−1^ to 0.5 s^−1^, our model shows that both clustering and ligand binding are promoted (Fig 4A-B). Using *E_II_* = 5 **k_B_T** and varying *E_IL_* from 3 *k_B_T* to 11 *k_B_T*, clustering is independent from *E_IL_* (Fig 4A), while overall ligand binding increases with *E_IL_* (Fig 4B). Clustering is mostly set by the strength of pairwise interactions between integrins, *E_II_*. It can be promoted by low *E_IL_*. and high *k_a_*, leading to a higher number of integrins able to diffuse and cluster (Fig 4A). Ligand binding is proportional to *E_IL_* at all *k_a_*. In experiments, variations in integrin activation rate are tied to variations in ligand binding affinity, making it unclear whether it is *k_a_* or *E_IL_* that determines organization of nascent adhesions. Our model shows that the rate of integrin activation set the level of the competition between ligand binding affinity and strength of pairwise interactions (Fig 4A).

**Fig 4.**
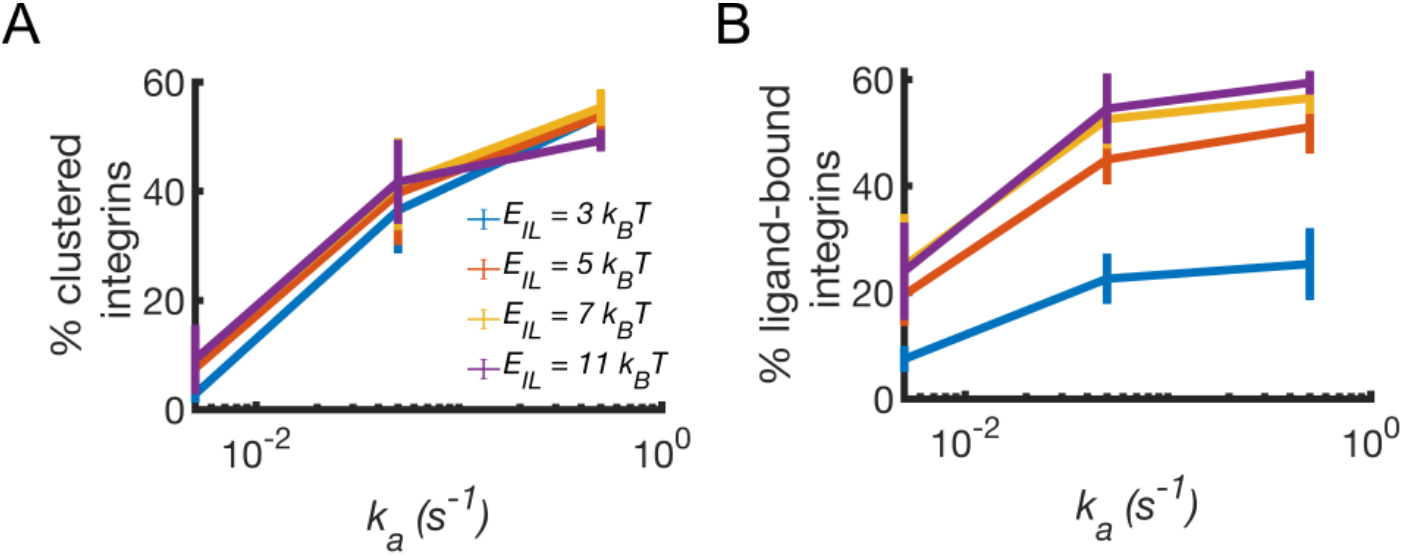
Integrin activation rate enhances clustering and ligand binding. (A) Average percentage of clustered integrins as a function of activation rate, *k_a_*, by varying ligand binding affinity, *E_IL_*, and keeping *E_II_* = 5 *k_B_T*. (B) Corresponding average percentage of ligand-bound integrins. Data are computed between 10-200 s of simulations from two independent runs. Error bars indicate standard deviation from the mean.

Experimentally, Mn^2+^ or antibodies are typically used to modulate ligand binding [55–58]. Both of these approaches, however, not only increase ligand binding affinity, but also the lifetime of the ligand bond. The increase of the ligand bond lifetime can be formally represented using a catch-bonds [59], where ligand unbinding rates decrease under tension and promotes stress transmission from the adhesions [60]. Therefore, we next used the model to test how variations in catch bond kinetics, combined with differences in the relative amount of β-1 and β-3 integrins, modulate ligand binding and stress transmission.

### Distribution of tension on integrins depends on bond dynamics

Since nascent adhesions transmit tension between the cytoskeleton and the ECM, we next asked how mixing integrins with different load-dependent bond kinetics impacts ligand binding and transmitted tension. The ß-1 and ß-3 integrins both behave as catch bonds that differ for unloaded and maximum lifetimes (Fig 2C) [49,53]. In the model, an increase in the percentage of ß-1 integrins while keeping the rest as ß-3 integrins, increases ligand binding from about 5% to 35% when using actin flow speeds below 15 nm/s (Fig 5A). The percentage of ligand-bound integrins is in direct proportion to the amount of ß-1 integrins (Fig 5A). At actin flow speeds below 15 nm/s, traction stress and flow rate are positively correlated, while at higher flows they are inversely correlated (Fig 5B), in agreement with previous findings [61,62]. Interestingly, variations in the relative fractions of the two integrin types do not affect the average tension on each integrin-ligand bond (Fig 5B). Below 10 nm/s actin flow, the minimum separation between ligand-bound integrins can be decreased from about 120 to 10 nm by increasing the fraction of β-1 integrins (Fig 4C). Stable adhesions, with minimum separation between ligand-bound integrins of 70nm, form with at least 20% ß-1 integrins (Fig 5C). Together, our results show that the relative fractions of β-1 and β-3 integrins cooperate with actin flow to determine ligand binding and adhesion stability.

**Fig 5.**
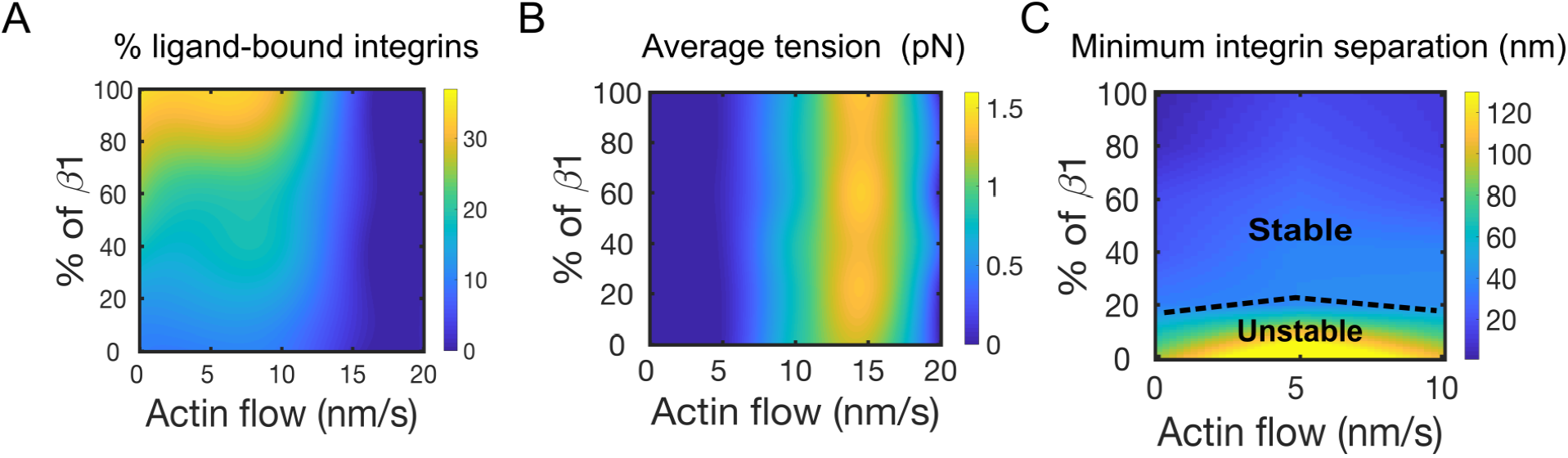
Amounts of β-1 and β-3 integrins determine ligand binding, traction stress, and adhesion stability. (A) Percentage of ligand-bound integrins by varying fraction of β-1 in a system of β-3 integrins, as a function of actin flow speed. (B) Average tension per integrin varying actin flow speed and percentage of simulated β-1 in a system of β-3 integrins. (C) Average distance between nearest ligand-bound integrins, without distinction between β-1 and β-3 integrins. Represented are regions of unstable and stable adhesions, depending upon the minimum spatial separation between any integrin type. Data are computed as averages between 1-120 s of simulations.

### Actin architecture does not impact integrin clustering and ligand binding, but changes the physical organization of integrins in adhesions

Interactions of adhesions with a cytoskeleton network play important roles in several cell activities, including spreading and migration [63]. The actin cytoskeleton exists in different architectures, depending on the cell location and function [64]. Therefore, we next consider how the architecture of the actin cytoskeleton can impact the formation of adhesions. We incorporate in the model explicit actin filaments, using random, crisscrossed, and bundled architectures (Fig 6A-C). The model assumes that ligand-bound integrins can interact with actin filaments, and that binding to actin increases integrin activation rate, as detected experimentally [65]. Increasing the fraction of β-1 integrins, ligand binding increases independent of network architecture (Fig 6D). By contrast, integrin clustering remains at about 20-30% when a percentage of β-3 integrins is used. When only β-1 integrins are used, integrin clustering decreases of about 3-fold, independent from network architecture (Fig 6E). The number of ligand-bound integrins with a separation less than 70 nm is enhanced using a bundled network architecture (Fig 6F). This suggests that the probability of adhesion stability and ultimately maturation is higher with bundled architecture relative to both crisscrossed and random distributions of actin filaments (Fig 6F).

**Fig 6.**
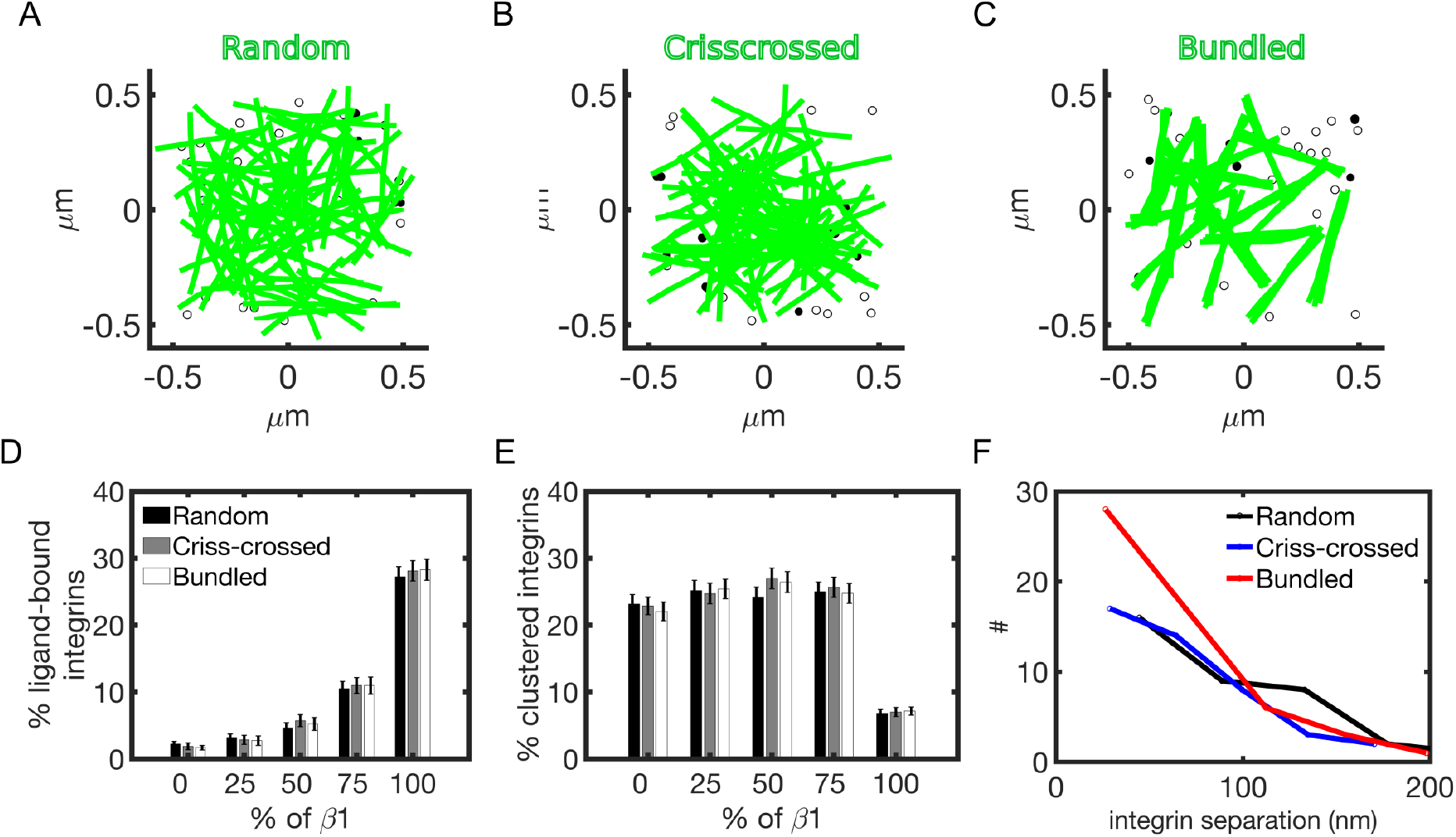
Actin architecture does not impact integrin clustering and ligand binding but changes the physical distribution of integrins with adhesions. (A) Snapshots from the simulations: random, crisscrossed and bundled actin networks above a layer of integrins. Filled circles indicate clustered integrins; empty circles indicate ligand-bound integrins. (B) Average percentage of ligand-bound integrins with respect to increasing amount of β-1 integrins (using *E_IL_* = *E_II_* = 9 *k_B_T*) at varying actin architectures. (C) Corresponding percentage of clustered integrins. (D) Distribution of average nearest neighbor distances between ligand-bound integrins using 50% β-1 integrins at varying actin architectures. Results are computed as averages between 20-30 s of simulation; error bars denote standard errors from the mean.

Collectively, our results indicate that the architecture of the actin cytoskeleton does not modulate the amount of ligand binding and integrin clustering. However, actin network architecture determines the physical distribution of ligand-bound integrins in adhesions with bundled actin filaments increasing the probability of adhesion stability, consistent with previous experimental observations [54,66].

## Discussion

Since different integrin types exist in nascent and mature adhesions (Fig 1B), a computational model is developed here in order to understand if differences in nanoscale physical properties of integrins reflect on adhesions. This is largely untested by experimental approaches because it is very challenging to simultaneously distinguish between integrin types and isolate their nanoscale physical properties. The model is used to study how ligand binding affinity, rate of integrin activation, strength of pairwise interactions, bond kinetics, as well as the architecture of a network of actin filaments modulate integrin organization in adhesions and stress transmission. Our results collectively show that ligand binding and integrin clustering are competing mechanisms and that a bundled actin networks favor adhesions stability, and ultimately maturation.

The model is developed through three consecutive stages of increasing complexity: (*i*) simulations of single-point integrins diffusing on a quasi-2D surface and switching between active and inactive states, binding ligands, and interacting laterally; (*ii*) incorporation of an implicit actin flow and integrin/ligand catch bonds kinetics; (*iii*) binding of integrins to semi-flexible actin filaments in either random, bundled, or crisscrossed architectures. At all stages, we distinguish between ß-1 and ß-3 integrins, by using either exact, experimentally detected physical parameters, realistic fold differences between the two, or estimates from previous free energy calculations.

For high *E_IL_*, many active integrins bind ligands and the fraction of integrins that can diffuse, and cluster, is reduced (Fig 3D-E). Accordingly, this happens when the fraction of ß-1 integrins is higher than that of ß-3 integrins (Fig 5A), since ß-1 integrins have higher ligand-binding affinity than ß-3 integrins [46]. By contrast, with many free diffusing integrins that have low *E_IL_*, and are less likely to bind ligands, the fraction of integrins that can encounter each other, and cluster is enhanced and reduces ligand binding (Fig 3D-E). This happens when the fraction of ß-3 integrins is higher than that of ß-1 integrins (Fig 5A) and can also depend on the higher diffusion coefficient of ß-3 integrins with respect to ß-1 integrins [45]. The result that ligand binding and integrin clustering are competing mechanisms is consistent with a kinetic Monte Carlo model showing that the thermodynamics of ligand binding and dynamics of integrin clustering interplay [48]. Our model reproduces this competing process over the same range of ligand binding affinities and strength of pairwise interactions. Moreover, previous studies in U2OS cells showed that ß-3 integrins cluster on both ß-3 and ß-1 integrin ligands, while ß-1 integrin clusters are present in adhesions only on ß-1 ligands [31]. Our result that ß-1 integrins are correlated with ligand-binding while ß-3 integrins are mostly responsible for clustering is consistent with the observation that clusters of ß-1 integrins are present only on ß-1 ligands, possibly because, in this case, ligand binding and not pairwise interactions facilitate adhesions assembly.

Our model also shows that the number of integrins per cluster, computed as the fraction of clustered versus ligand bound integrins, is in the range of 2-15 particles, depending on *E_II_* and *E_IL_* (Fig 3E). This value is comparable to the experimentally estimated number of integrins in nascent adhesions, between 5-7 [10]. By varying ligand density in the model, the ratio between clustered and ligand bound integrin particles does not vary significantly (Fig S1), suggesting that the average number of integrins per cluster in nascent adhesions is not modulated by ligand concentration, consistent with previous experimental observations [10].

In the presence of actin flow, the fraction of ß-1 integrins is positively correlated with ligand binding (Fig 5A). Above a threshold actin flow, however, ligand binding is almost suppressed, independent from relative amounts of ß-1 and ß −3 integrins (Fig 5A), because of faster ligand unbinding from both integrins. This reduction in bound ligands corresponds to a drop in the average tension per integrin upon increasing actin flow (Fig 5B). The biphasic response of tension to actin flow was previously observed experimentally [61] and is consistent with models of adhesion clutch assembly and rigidity sensing [67]. By increasing the fraction of ß-1 integrins, a reduction of lateral integrin spacing is observed with our model (Fig 5C). Previous studies on the lateral separation of integrins in adhesions reported that a minimum spacing of 70 nm is required to form stable adhesions [68]. This value corresponds in the model to a minimum of 20% ß-1 integrins (Fig 5B). This value represents a prediction from our CG model that can be experimentally tested in the future. When only ß-3 integrins are used in the model, their lateral separation, upon binding ligands, is about 120 nm (Fig 5C), supporting the notion that ß-1 integrins are needed to form stable adhesions. This is consistent with the fast binding/unbinding dynamics of ß-3 integrins previously observed in experiments [25].

By incorporating a positive feedback between actin filament engagement and integrin activation, as observed in [65,69], the competition between clustering and ligand binding is maintained in all actin architectures (Fig 6D-E). The number of ligand-bound integrins with an average separation below 70 nm is enhanced with a bundled architecture (Fig 6F), suggesting that this configuration favors adhesion stability, and ultimately maturation [54]. Of interest for future studies is mimicking conditions of actin filament turnover and dynamic ligands, in order to understand how a dynamic cytoskeleton can interplay with integrin mixing in forming nascent adhesions on dynamic substrates.

To conclude, with our highly coarse-grained model based on Brownian Dynamics, we extend the scope of previous theoretical and computational studies of integrin-based adhesions formation, by testing how differences in nanoscale properties of ß-1 and ß-3 integrins impact ligand binding, clustering and transmission of traction stress. By coupling physical parameters (such as diffusivity) together with chemical (i.e., affinity and receptor pairwise interactions) and mechanical (bond kinetics) parameters, and by using an explicit actin cytoskeleton, our model shows that nascent adhesions assembly is finely tuned by differences in nanoscale physical properties of integrins. The CG model ultimately demonstrates that nanoscale differences in integrin dynamics are sufficient to determine ligand binding and integrin clustering. By incorporating dynamics of individual integrins in an explicit way, our model provides results that are consistent with a number of previous independent experimental observations, revealing important insight into the molecular origins of adhesion organization and mechanics. Taken together, our modeling results support the general view that a cell can control integrin expression to determine morphological and dynamic properties of adhesions.

## Methods

In order to characterize how nanoscale physical properties of integrins impact the assembly of nascent adhesions, we developed a highly coarse-grained computational model based on Brownian Dynamics. The model is agent-based in the way sequential dependencies regulate interactions between integrin/ligand, integrin/integrin, and integrin/actin. Integrins, ligands, and actin filaments are explicit particles, while the cell membrane is implicit. Solvent molecules mimicking cytoplasmic effects are replaced by stochastic forces, depending on cytoplasmic viscosity. Inactive integrins diffuse and, when active, can bind ligands and interact laterally. When integrins are bound to ligands, they can engage actin filaments. The interaction between integrins and actin filaments locally increases integrin activation rate, ultimately resulting in a positive feedback between actin binding and ligand binding [65,69].

In order to distinguish between β-1 and β-3 integrins, we examine the effect of varying integrin activation rates, motility, ligand binding affinity, clustering, and bond kinetics (Fig 2A-C). By varying the relative amounts of β-1 and β-3 integrins, we analyze fractions of ligand-bound integrins, clustered integrins, and average tension on integrin-ligand bonds (Fig 3–5). Moreover, we study the effect of different actin filaments architecture on adhesions morphology (Fig 6).

The model is an extension of our mechanosensing model [5] but differs from it in several ways. First, each integrin exists in either active or inactive state, determined by activation and deactivation rates. Second, the model incorporates tunable parameters for integrin physical properties, allowing us to discriminate between integrin types. Third, explicit semiflexible actin filaments are included.

### Computational domain

The computational domain includes two main systems: a square bottom surface, of 1 μm per side, and a rectangular 3D domain above the surface, with dimensions 1 x 1 x 0.04 μm (Fig 2A). The square bottom surface mimics the substrate; the bottom side of rectangular domain mimics the ventral cell membrane above the substrate, while its inside space represents a 40 nm thick cytoplasmic region where actin filaments diffuse beyond the ventral membrane (Fig 2A). The cell membrane is separated from the substrate by 20 nm, a dimension characteristic of active integrin headpiece extension (Fig 1A) [70]. Within the cell membrane, integrins diffuse in quasi-2D and are restrained in the vertical direction by a weak harmonic potential with spring constant 100 pN/μm, mimicking membrane friction. In the cytoplasmic region, a repulsive boundary is used on the top surface, to avoid filaments crossing the boundary. Periodic boundary conditions are applied on all sides of the domain, in order to avoid finite size effects.

### Integrin and ligands representation

The model considers a given number of ligands on the substrate, randomly distributed and fixed in space. We use a ligand density of 1000#/μm^2^, of the same order of that used in a previous model of adhesions assembly [71]. Integrin density on the cell membrane is ~100#/μm^2^ [5]. Integrins are initially randomly distributed and diffuse over the course of the simulations, with diffusion coefficient *D* = 0.1 μm^2^/s for ß-1 integrins and *D* = 0.3 μm^2^/s for ß-3 integrins [45]. Introducing volume exclusion effects between integrins, in the form of a weak repulsion between nearby particles (1 pN force), does not change the fraction of ligand-bound integrins, their average separation, the mean tension per integrin and its distribution (see Fig S2, A-D). Increasing the magnitude of this repulsion (10 pN), however, affect the average separation of integrins (Fig S2 E-F).

### Actin filament representation

Semiflexible polymers represent actin filaments as spherical particles connected by harmonic interactions. Filaments have fixed length of 0.5 μm, corresponding to 6 beads separated by 0.1 μm equilibrium distance. The model of actin filaments is explained in detail in [72]. Actin filament beads are subjected to both stochastic and deterministic forces. Stochastic forces on the i-th bead are random in direction and magnitude in order to mimic thermal fluctuations and satisfy the fluctuation-dissipation theorem:

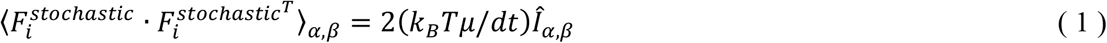

with *Î_α,β_* being the second-order unit tensor [62] and *μ* being a friction coefficient equal in three directions.

Deterministic contributions come from bending and extensional forces on the filament beads. The bending force is computed as:

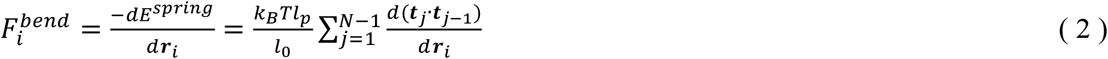

where *l_p_* = 10 μm is actin filament the persistence length, *N* is the number of beads in a filament (*N* = 6) and 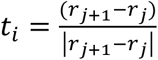.

The extensional force on filaments beads is computed as:

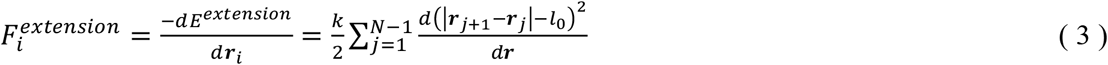

where *l*_0_ is the equilibrium length of 0.1 μm, *l*_0_ is the spring constant of 100 pN/μm.

Each spherical particle of a filament represents a binding site for integrin and each binding site can interact with multiple integrins.

### Agent-based algorithm

In order to mimic hierarchical formation of nascent adhesions [10], the algorithm incorporates sequential interactions between integrins, ligands and actin filaments. First, we simulate a system composed of only integrins and ligands, in order to explore the ways in which integrins cluster and bind ligands in an actin-independent way. Then, we add actin filaments and study the effect of actin network architecture on adhesions formation.

Integrins switch between inactive and active states, with rates of activation and deactivation *k_a_* = 0.5 s^−1^ and *k_d_* = 0.0001 s^−1^, of the same orders of those previously estimated [46,47,71]. Activation probability corresponds to *P_a_* = *k_a_dt*, with time-step *dt* = 0.0001 s. Upon activation, integrins can interact with free ligands, using a harmonic potential (with equilibrium separation 20 nm and spring constant 1 pN/μm), and cluster with other active integrins, depending on relative distances. Ligand binding occurs within a threshold distance of 20 nm, which reflects the extension of the open conformation of αпbβ_3_ integrin away from the membrane [70]. Each integrin can bind only one ligand, and each ligand can bind only one integrin, mimicking binding sites specificity. Clustering occurs below a threshold of 30 nm, a value of the same order of the integrin-to-integrin lateral separation observed experimentally [70] and one order of magnitude lower than the minimum separation between individual adhesions [73].

The probability of integrin deactivation is *P_d_* = *k_d_dt*. Once inactive, integrin loses its connections with ligands and other integrins. Integrins unbind ligands with dissociation probabilities depending on their affinities: *P* = *e^E_lL_^dt*. They break later connections with probabilities inversely proportional to strength of pairwise interaction: *P* = *e^E_II_^dt*. For ß-1 integrins, we use high affinity, ~9 *k_B_T*; for ß-3 integrins we use low affinity, 3-5 *k_B_T*.

Ligand-bound integrins can establish harmonic interactions with semiflexible actin filaments below 5 nm, approximating the size of the intracellular integrin tails [70]. Interaction between integrin and actin is harmonic with equilibrium distance of 3 nm and spring constant of 1 pN/μm [70]. This interaction simplifies a layer of adhesion molecules, including vinculin, talin and α-actinin.

### Brownian Dynamics simulations via the Langevin Equation

Displacements of integrin and actin filament particles are governed by the Langevin equation of motion in the limit of high friction, thus neglecting inertia:

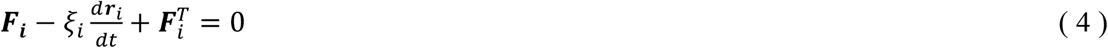

where **r**_*i*_ is a position vector of the *i*th element, *ζ_i_* is a drag coefficient equal in three directions, *t* is time, **F**_*i*_ is a deterministic force, and **F**_*i*_^T^ is a stochastic force satisfying the fluctuation-dissipation theorem [74]. **F**_*i*_ is the sum of forces resulting from interactions of integrins with a ligand and/or other particles in the system, and actin flow in a direction parallel to the substrate Positions of the various elements are updated at every time step using explicit Euler integration scheme:

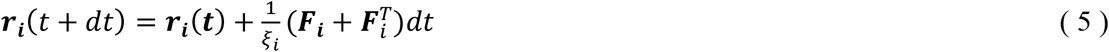

### Integrin/ligand unbinding formalism

Since contraction forces are not needed for the assembly of nascent adhesions [5,10,66], our computational model only incorporates forces mimicking actin retrograde flow. In order to simulate actin flow and characterize distribution of traction stress at various flow rates, a constant force is applied on ligand bound integrins, along *y* (Fig 2B). Lifetime of the bond between integrin and ligand follows the catch-bond formalism (Fig 2C), using: for ß-1 integrins an unloaded affinity of 2 s and a maximum lifetime of 15 s; for ß-3 integrins an unloaded affinity of 0.5 s and a maximum lifetime of 3 s. The parameters for the catch bond kinetics are from previous experimental characterizations [49,53,60]. Curves of bond lifetime versus tension are shown in Fig 2C.

For ß-1 integrins, we implemented an unbinding rate as a function of the force acting on the bond, *F*:

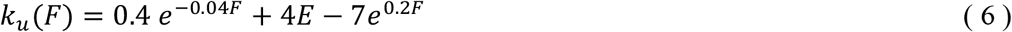

For ß-3 integrins, we used unbinding rate:

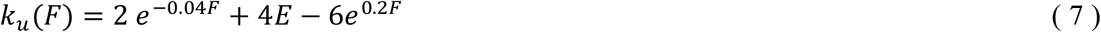

### Positive feedback between filament binding and integrin activation

To mimic promotion of integrin clustering upon ligand binding and actin filament engagement [65], we introduce a positive feedback between binding of integrin to a filament and integrin activation rate. In the model, integrins can bind a filament only if already bound to a ligand particle. Upon binding to actin, integrin activation rate is increased by 2 to 4% relative to its initial value. This assumption is motivated by recent evidence from TIRF experiments on T-cells, where it was demonstrated that actin binding and correct ligand positioning are needed for integrin activation [65]. We use the model with the positive feedback (schematics in Fig 2D) to test the effect of different actin architectures on ligand binding and clustering (Fig 6).

### Experimental approach

Human Foreskin Fibroblasts (HFF) were purchased from ATCC and cultured in DMEM media (Mediatech) supplemented with 10% Fetal Bovine Serum (Corning), 2 mM L-glutamine (Invitrogen) and penicillin-streptomycin (Invitrogen). HFFs were plated on glass coverslips incubated with 10 μg/mL fibronectin (EMD Millipore) for 1 hr at room temperature. Cells were fixed 1 hr after plating by rinsing them in cytoskeleton buffer (10 mM MES, 3 mM MgCl2, 1.38 M KCl and 20 mM EGTA) and then fixed, blocked and permeabilized in 4% PFA (Electron Microscopy Sciences), 1.5% BSA (Fisher Scientific), and 0.5% Triton X-100 (Fisher Scientific) in cytoskeleton buffer at 37° for 10 minutes. Coverslips were subsequently rinsed three times in PBS and incubated with either a β_1_ antibody (1:100; Abcam product #:ab30394) or β_3_ antibody (1:100; Abcam product #:ab7166) followed by AlexaFluor 488 phalloidin (1:1000; Invitrogen) and a AlexaFluor647 donkey anti-mouse secondary antibody (1:200; Invitrogen).

Cells were imaged using a 1.2 NA 60X Plan Apo water immersion lens on an inverted Nikon Ti-Eclipse microscope using an Andor Dragonfly spinning disk confocal system and a Zyla 4.2 sCMOS camera. The microscope was controlled using Andor’s Fusion software.

## Acknowledgements

This research was supported by the Department of Defense Army Research Office through MURI grant W911NF1410403. It was also partially supported by the University Chicago Materials Research Science and Engineering Center, which is funded by the National Science Foundation under award number DMR-1420709. Computer time was provided by the University of Utah Research Computing Center.

## Supporting information

**S1 Fig.**
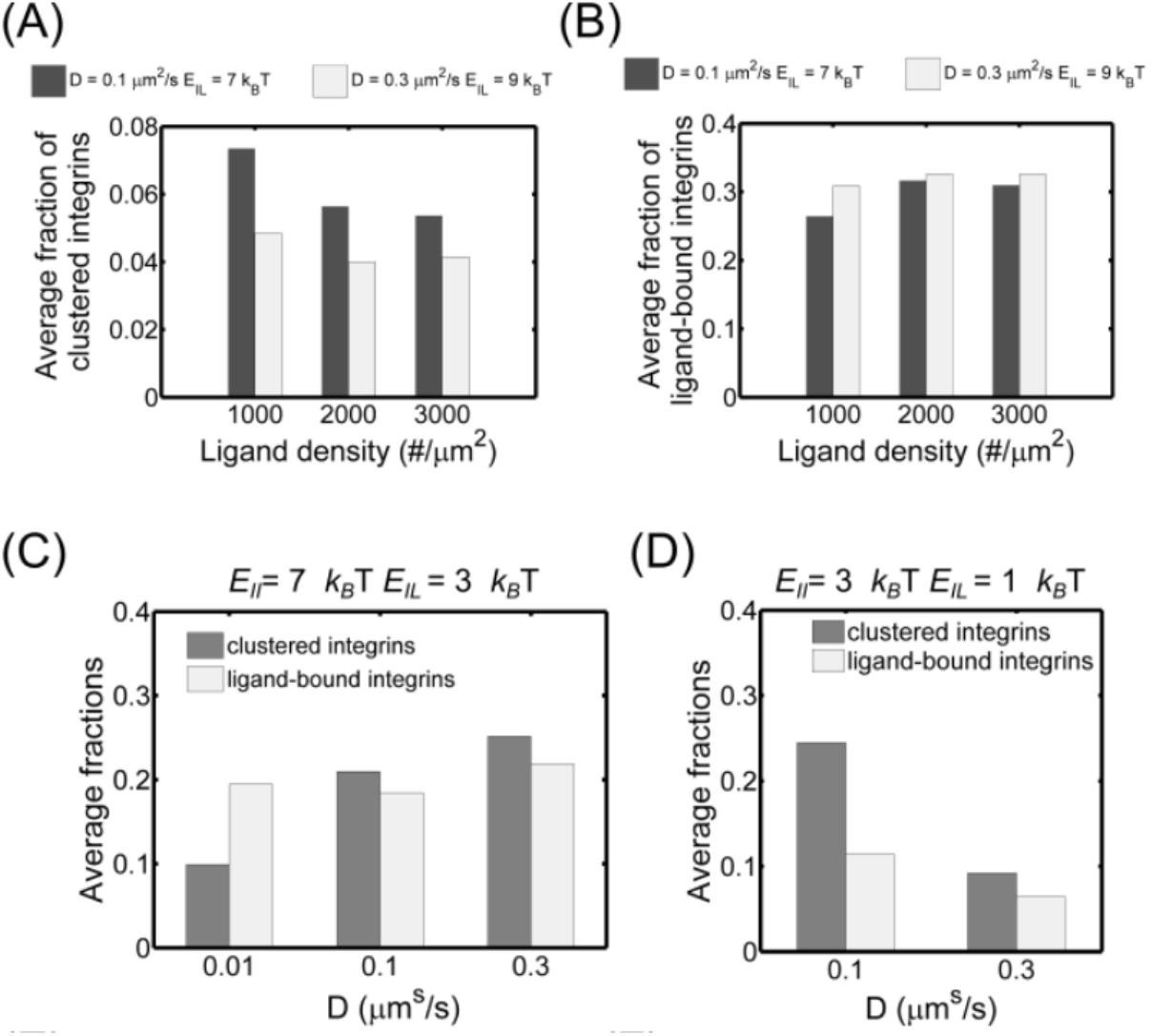
Integrin diffusivity cooperate with ligand density, strength of pairwise interactions, and affinity to mediate clustering and ligand binding. (A) Average fraction of clustered integrins and (B) corresponding fraction of ligand-bound integrins varying ligand density for two conditions of integrin properties: *D* = 0.1 μm^2^/s, *E_IL_* = 7 *k_B_T*, *E_II_* = 9 *k_B_T* ; *D* = 0.3 μm^2^/s, *E_IL_*= 9 *k_B_T*, *E_II_*= 9 *k_B_T*. Data are computed between 80-100 s of simulations from three independent runs. (C) Average fractions of clustered and ligand-bound integrins varying diffusion coefficient, using *E_IL_* = 3 *k_B_T* and *E_II_* = 7 *k_B_T*. Data are computed between 200-600 s of simulations, from three independent runs. (D) Average fractions of clustered and ligand-bound integrins at *D* = 0.1 and 0.3 μm^2^/s, *E_IL_* = 1 *k_B_T* and *E_II_* = 3 *k_B_T*. Data are computed between 200-600 s of simulations, from three independent runs.

**S2 Fig.**
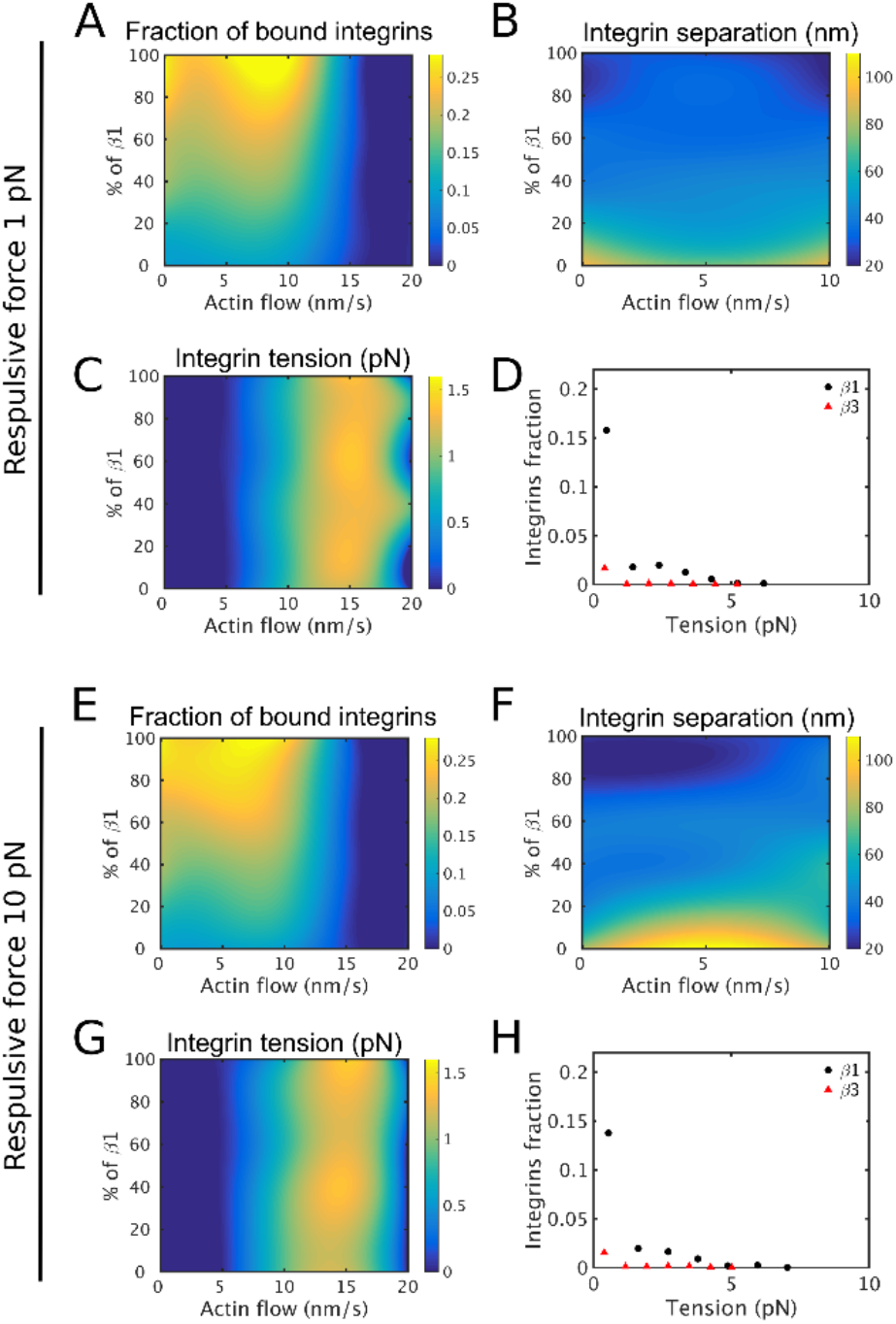
Volume exclusion effects between integrins do not change ligand binding and traction stress as a function of actin flow. (A) Average fraction of ligand-bound integrins as a function of actin flow speed and percentage of β-1 integrins in a system of β-3 integrins. (B) Corresponding average nearest neighbor distance between ligand-bound integrins. (C) Corresponding average tension per integrin. (D) Distribution of tension on ligand-bound integrins for the two integrins types, using 80% β-1 and 20% β-3 integrins and 10 nm/s actin flow. Data are computed between 1-20 s of simulations, using a weak repulsive potential (1 pN) between integrins closer than 1 nm. Panels E-H show data as in panels A-D for implemented repulsive forces of 10 pN.

